# CAP-MAP: Cap Analysis Protocol with Minimal Analyte Processing, a rapid and sensitive approach to analysing mRNA cap structures

**DOI:** 10.1101/2019.12.29.890368

**Authors:** Alison Galloway, Abdelmadjid Atrih, Renata Grzela, Edward Darzynkiewicz, Michael A. J. Ferguson, Victoria H. Cowling

## Abstract

Eukaryotic messenger RNA (mRNA) is modified by the addition of an inverted guanosine cap to the triphosphate at the 5’ end. The cap guanosine and initial transcribed nucleotides are further methylated by a series of cap methyltransferases to generate the mature cap structures which protect RNA from degradation and recruit proteins involved in RNA processing and translation. Research demonstrating that the cap methyltransferases are regulated has generated interest in determining the methylation status of the mRNA cap structures present in cells. Here we present CAP-MAP: Cap Analysis Protocol with Minimal Analyte Processing, a rapid and sensitive method for detecting cap structures present in mRNA isolated from tissues or cell cultures.

## Introduction

Eukaryotic RNAPII (RNA polymerase II)-transcribed RNAs are modified by the addition of 7-methylguanosine to the triphosphate found on the 5’ first transcribed nucleotide, forming the cap structure denoted ^m7^GpppN, (N is any nucleotide) (Fig.1) [1,2]. For many short RNAPII transcripts involved in guiding RNA processing and modification the cap is a precursor for further modification [3]. For pre-messenger RNA (mRNA), the cap structure guides transcript processing and selection for translation through interactions with cap binding proteins, whilst protecting the transcript from 5’-3’ exonucleases [4]. Metazoan mRNA caps additionally contain ribose methylated on the O-2 position (denoted N_m_) on the first and second transcribed nucleotides which creates a further “self mRNA” mark, enabling innate immunity proteins to differentiate it from unmethylated, foreign RNA [5,6]. Removal of the cap by decapping enzymes usually directs mRNA to be degraded. Decapping is also regulated by RNA binding proteins which promote or antagonise the binding of decapping complexes to the mRNA [7]. Thus, the cap is essential for the proper processing, function and lifespan of a mRNA.

**Figure 1.**
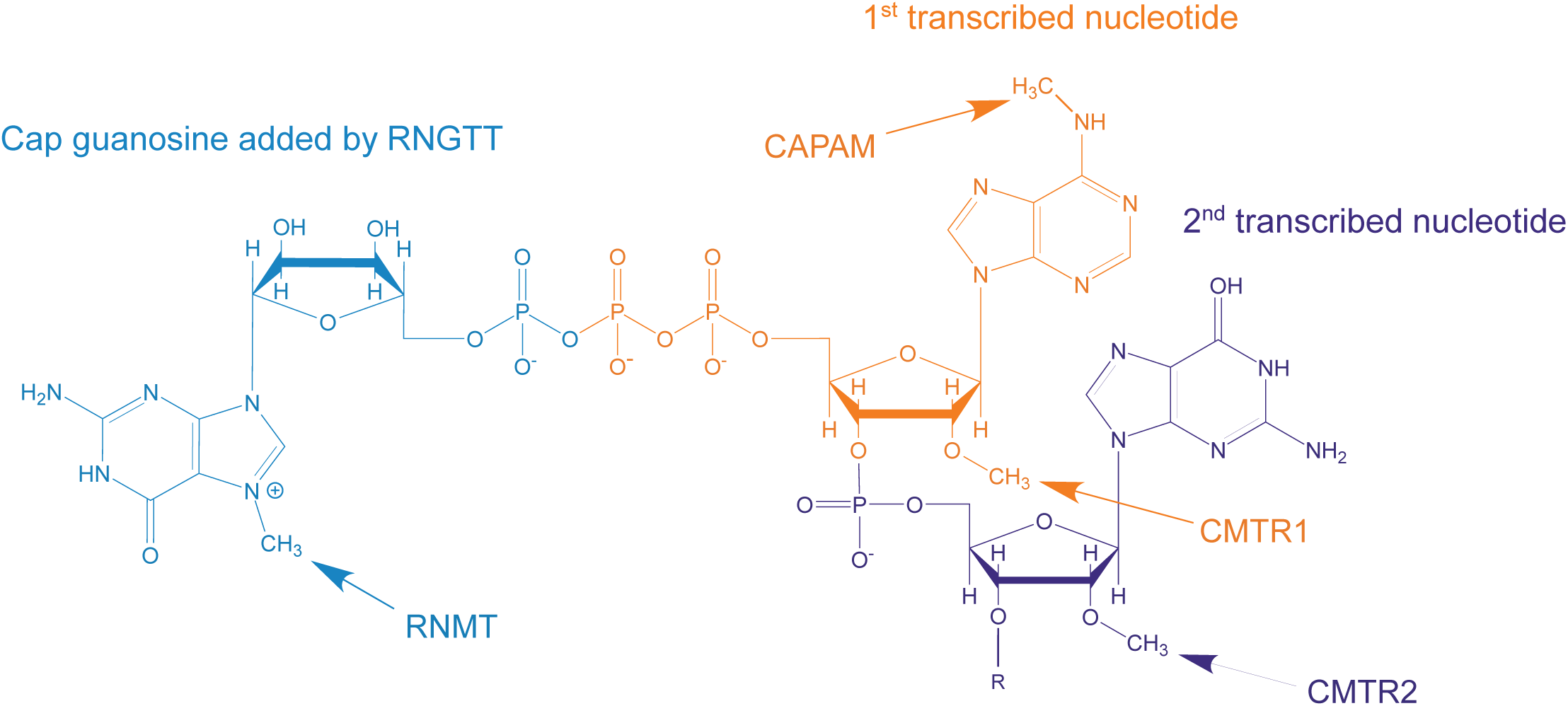
mRNA cap structure. A common cap structure is depicted, including cap guanosine, first transcribed nucleotide and second transcribed nucleotide. The sites of action of the capping enzymes RNGTT, RNMT, CMTR1, CMTR2 and CAPAM are indicated.

Mammalian mRNA processing initiates co-transcriptionally with the addition of the inverted guanosine cap, catalysed by Capping Enzyme/RNA guanylyltransferase and 5’ phosphatase (CE/RNGTT) [2,8]. The terminal cap guanosine is methylated by RNA cap methyltransferase (RNMT) on the N-7 position and the first two transcribed nucleotides are modified by cap-specific methyltransferases as follows: the O-2 position of the ribose of the first and second transcribed nucleotides are methylated by Cap-Methyltransferase 1 (CMTR1) and CMTR2 respectively, and if the first transcribed nucleotide is adenosine it is methylated on the N-6 position by Cap-specific Adenosine Methyltransferase (CAPAM, also known as PCIF1) [9-12] (Fig.1).

The cap structure was first elucidated in viral mRNA and determined to mimic eukaryotic cellular mRNA in order to hijack the translation machinery and evade detection by the host innate immunity proteins [13]. In the 1970s, ^m7^GpppN_m_pN and ^m7^GpppN_m_pN_m_ caps, as well as the ^m6^A_m_ modification were detected in mammalian cells by chromatographically separating RNAse-digested radiolabelled mRNA and by mass spectrometry [14-20]. More recently it is has become apparent that the cap methyltransferases are regulated during important cellular processes including the cell cycle, oncogenic transformation, anti-viral responses and embryonic stem cell differentiation [21-27]. This has rejuvenated interest in determining the relative abundance of each cap structure and additional cap analysis techniques have been developed [28]. First nucleotide ribose O-2 methylation and adenosine N-6 methylation have been detected by radiolabelling the terminal nucleotides and resolving them by 2D TLC [29]. Since this method does not require *in vivo* labelling, mRNA from any source can be analysed and the method can be adapted to investigate methylation of specific mRNAs, however, it is technically challenging and requires the use of radio-isotopes. Affinity reagents can be used to enrich methylated nucleotides and specific cap structures in a semi-quantitative manner. Capped mRNA can be enriched using antibodies that recognise ^m2,2,7^G and ^m7^G, although such antibodies exhibit gene-specificity which may be due to the cap structures present or RNA secondary structure [25,30,31]. Recent studies have detected internal ^m7^G in mRNA, which compromises the use of anti-7-methylguanosine antibodies for cap detection [32,33]. Similarly internal and cap adjacent ^m6^A have been mapped using an anti-^m6^A antibody, but since internal ^m6^A is abundant this method requires knowledge of the transcription start site to determine ^m6^A-containing cap structures. [34,35]. Recombinant eIF4E can also be used as an affinity reagent for capped mRNA although this does not specify particular cap structures [4]. Recently, an antibody-free ^m6^A mapping method has been developed [36]. A cytidine deaminase fused to the ^m6^A-binding domain of YTH domain was expressed in cells, resulting in C-to-U deamination at sites adjacent to ^m6^A residues which can be detected by RNA-seq. With the advent of nanopore sequencing, methylated nucleotides can be detected on specific sequences and this is likely to include the first transcribed nucleotides of the cap in the near future [37].

Mass spectrometry has been employed to detect various nucleotide modifications, including the cap [1,38,39]. Since the first transcribed nucleotide is expected to be N_m_ (ribose O-2 methylated), the presence of ^m6^A_m_ is associated with cap structures. A more accurate method for cap analysis involves detecting the cap dinucleotide ^m7^GpppN_m,_ or short 5’ mRNAs including this structure. Recently, this has been used successfully to detect cap structures derived from short RNAs, precursor tRNAs, as well as to monitor the activities of CAPAM and the ^m6^A demethylase fat mass and obesity-associated protein (FTO) on the first transcribed nucleotide of *in vitro* transcribed RNA [3,26,40-42]. More recently, cellular mRNA cap structures isolated with an anti-^m7^G antibody have been analysed by mass spectrometry and ^m7^GpppG_m_G_p_, ^m7^GpppA_m_G_p_ and ^m7^Gppp^m6^A_m_G_p_ caps were readily detected [9]. Caps lacking first transcribed nucleotide methylation were not detected indicating they are uncommon.

To gain a better understanding of *in vivo* cap regulation we developed an approach to determine the relative proportions of the different mRNA cap structures using a rapid and unbiased approach. Here we present the CAP-MAP (Cap Analysis Protocol with Minimal Analyte Processing) method for detecting and quantifying mRNA cap structures by liquid chromatography–mass spectrometry (LC-MS).

## Results

We developed CAP-MAP (Cap Analysis Protocol with Minimal Analyte Processing) to detect and quantitate the mRNA cap structures present in cells and tissues. Specifically, the method can assess the permutations of N-7 methylation of the terminal cap guanosine, O-2 methylation of the first nucleotide ribose and N-6 methylation of first nucleotide adenosine in mRNA caps from biological samples (Fig.1). Briefly, the method involves mRNA enrichment from total cellular RNA using oligo-dT affinity beads. The mRNA was then digested with nuclease P1, a non-specific ssRNA/DNA nuclease, to release nucleotide 5’-monophosphates and cap dinucleotides, ^(m7)^GpppN_(m)_. The latter were resolved by liquid chromatography on a porous graphitic carbon (PGC) column and identified by negative ion electrospray mass spectrometry using multiple reaction monitoring (MRM) (Fig.2). Negative ion mode was selected because of the propensity of nucleotides to form negative ions and for its better signal : noise ratio relative to positive ion mode. MRM is a procedure whereby, at any given moment, the first quadrupole/mass filter is programmed to allow through ions of only a single *m/z* value (X) into the second quadrupole/collision cell, where collision induced dissociation (CID) occurs, and the third quadruple/mass filter is programmed to allow through ions of typically only two *m/z* values (Y and Z) to the detector. In this way, ion currents (peaks) are only recorded when an analyte that produces a precursor ion (in this case an [M-H]^-^ ion) of *m/z* X generates product ions of *m/z* Y and Z. This, together with the LC retention time, provides confidence in the analyte identification as well as great sensitivity since the mass spectrometer is not spending time scanning irrelevant mass ranges.

**Figure 2.**
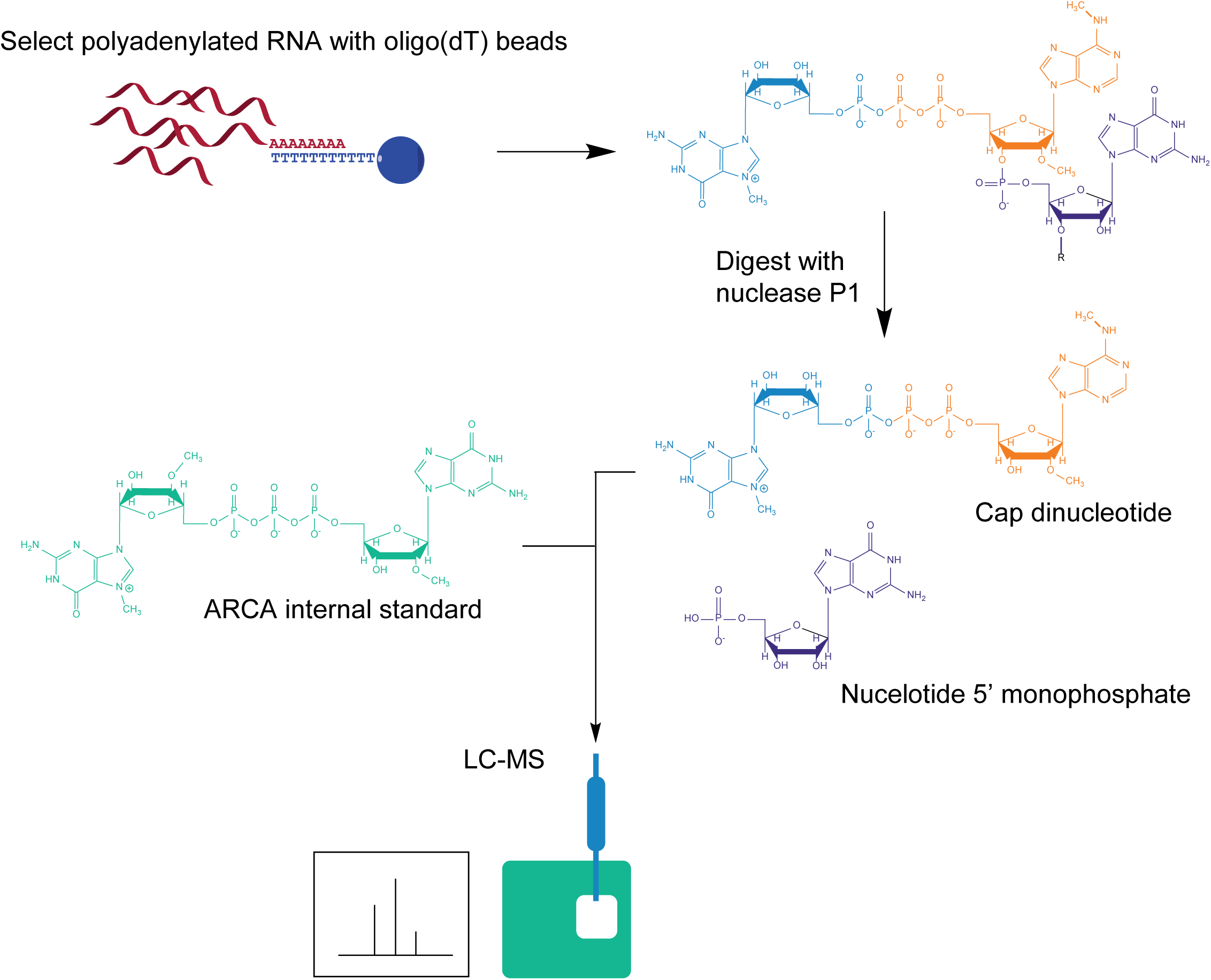
Overview of RNA preparation for CAP-MAP analysis. Cellular RNA is purified on oligo (dT) conjugated beads and digested with P1 nuclease to release cap dinucleotides and nucleotide monophosphates. The synthetic cap standard, ARCA (anti-reverse cap analogue), is added to the digested nucleotides. The sample is run on a PGC (porous graphitic carbon) column coupled to a triple quadrupole mass spectrometer operating in negative ion mode and programmed to detect cap dinucleotides in the multiple reaction monitoring (MRM) mode.

### LC-MS analysis of cap dinucleotides

Eleven synthetic cap dinucleotides, four obtained commercially from NEB and seven custom-synthesised (Table 1), were used to established retention times relative to an internal standard on PGC liquid chromatography and to optimise multiple reaction monitoring (MRM) conditions (mass transitions and collision energies) for their detection and quantification (Table 2). The internal standard was the so-called “anti-reverse cap analogue” (ARCA), which is not present in cells. ARCA has the structure ^m7^G_O-3m_pppG, which is physically similar but structurally distinct from endogenous cellular caps. During method development, we selected HyperCarb PGC as the preferred LC column type over SeQuant ZICpHILIC because the latter could not resolve the isobaric cap structures ^m7^Gppp^m6^A and ^m7^GpppA_m_. Using PGC, we found that excellent peak-shape and low carry-over between runs (<0.1%) could be obtained by using a pH 9.15 aqueous component and maintaining the column at 45°C. Further, the high organic solvent (acetonitrile) content required for cap elution from PGC was advantageous in the electrospray ionisation process. Using the conditions described in materials and methods, all eleven cap dinucleotides could be resolved by retention time and/or MRM transitions (Fig.3). To reduce PGC column deterioration over time, leading to peak-broadening and increasing retention times (West et al., 2010; Tashinga et al. 2016), we adopted some of the recommendations of (Tashinga et al., 2016 and Jansen et al. 2009). In particular, we regularly regenerated the columns by equilibrating in 95% methanol for between 1 h and 16 h.

**Figure 3.**
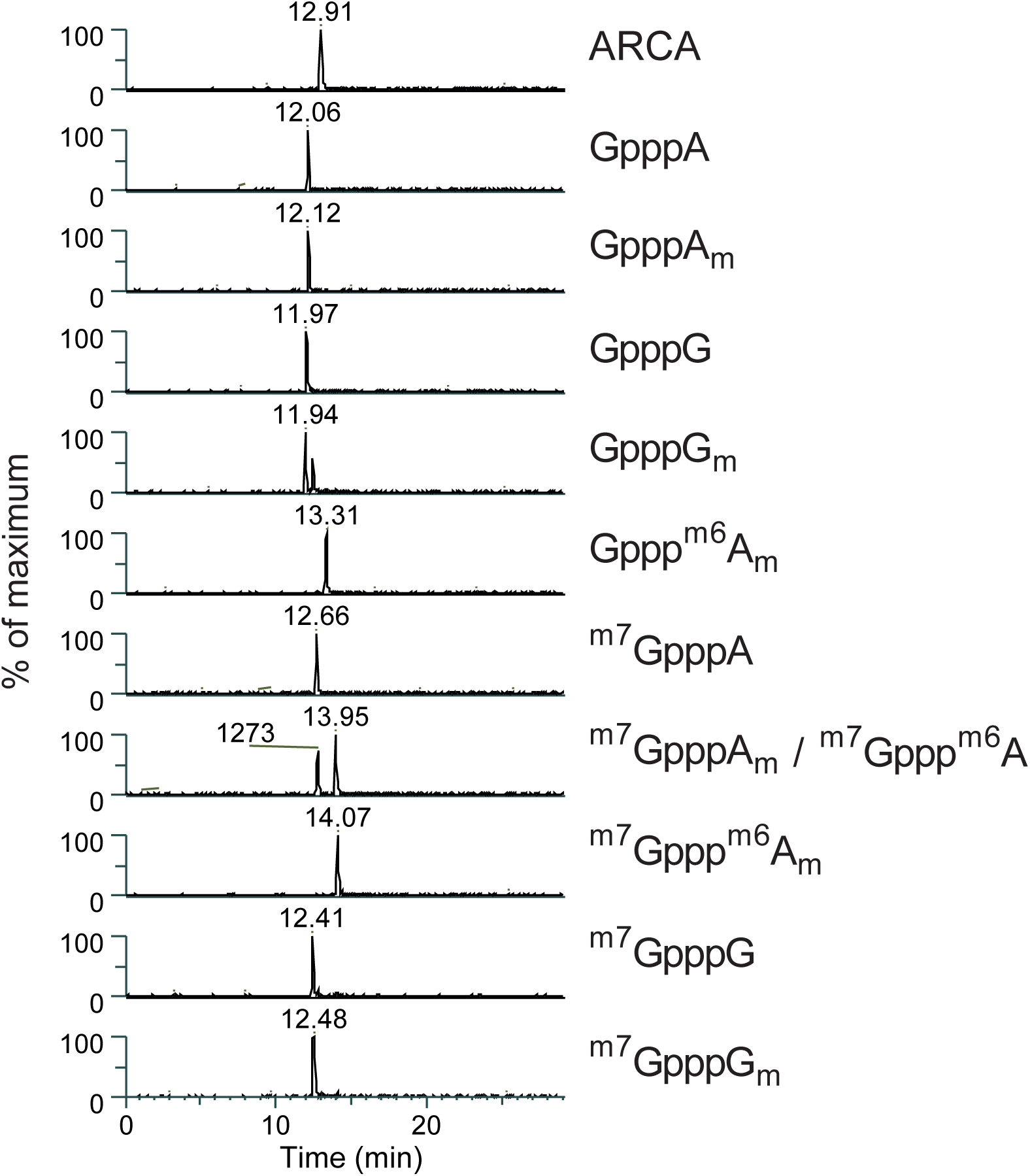
Elution profiles of 11 cap nucleotides on a PGC column. Chromatogram showing the differential separation of 11 cap nucleotides on a PGC (porous graphitic carbon) column. In the lane containing ^m7^GpppA_m_ and ^m7^Gppp^m6^A the ^m7^GpppA_m_ elutes earliest.

To determine the linear range of detection for the cap dinucleotides during LC-MS a dilution series of ten of the cap dinucleotides was analysed (Fig.4 A, B). All cap dinucleotides were detected in a linear range down to ∼4 fmol, with detection of some caps being non-linear at lower concentrations. Linear regression was used to calculate the slope for the peak area/fmol of each cap which was later used to convert peak area to fmol quantities for each cap (Table 3).

^m7^GpppG and GpppG_m_ caps are a challenge to distinguish by LC-MS because they are isobaric, elute very closely on PGC and produce similar CID spectra. Fortunately, ^m7^GpppG generates a unique product ion at *m/z* 635.9; thus we can detect and quantify ^m7^GpppG with the unique precursor -> product ion transition of *m/z* 800.9 -> 635.9. However, both ^m7^GpppG and GpppG_m_ produce precursor -> product ion transitions of *m/z* 800.9 -> 423.9 and 438.0. Thus, to quantify GpppG_m_ we have to correct for any *m/z* 800.9 -> 423.9 and 438.0 transition ion currents due to ^m7^GpppG. This is possible because the ratio of *m/z* 635.9 : 423.9 : 438 product ion intensities for ^m7^GpppG are constant. Consequently, the *m/z* 423.9 and 438 product ion contributions due to ^m7^GpppG can be back-calculated from the ^m7^GpppG-unique *m/z* 635.9 ion current. Examples of the calculations of the concentrations of ^m7^GpppG and GpppG_m_ in liver samples are shown in Fig.4E.

**Figure 4.**
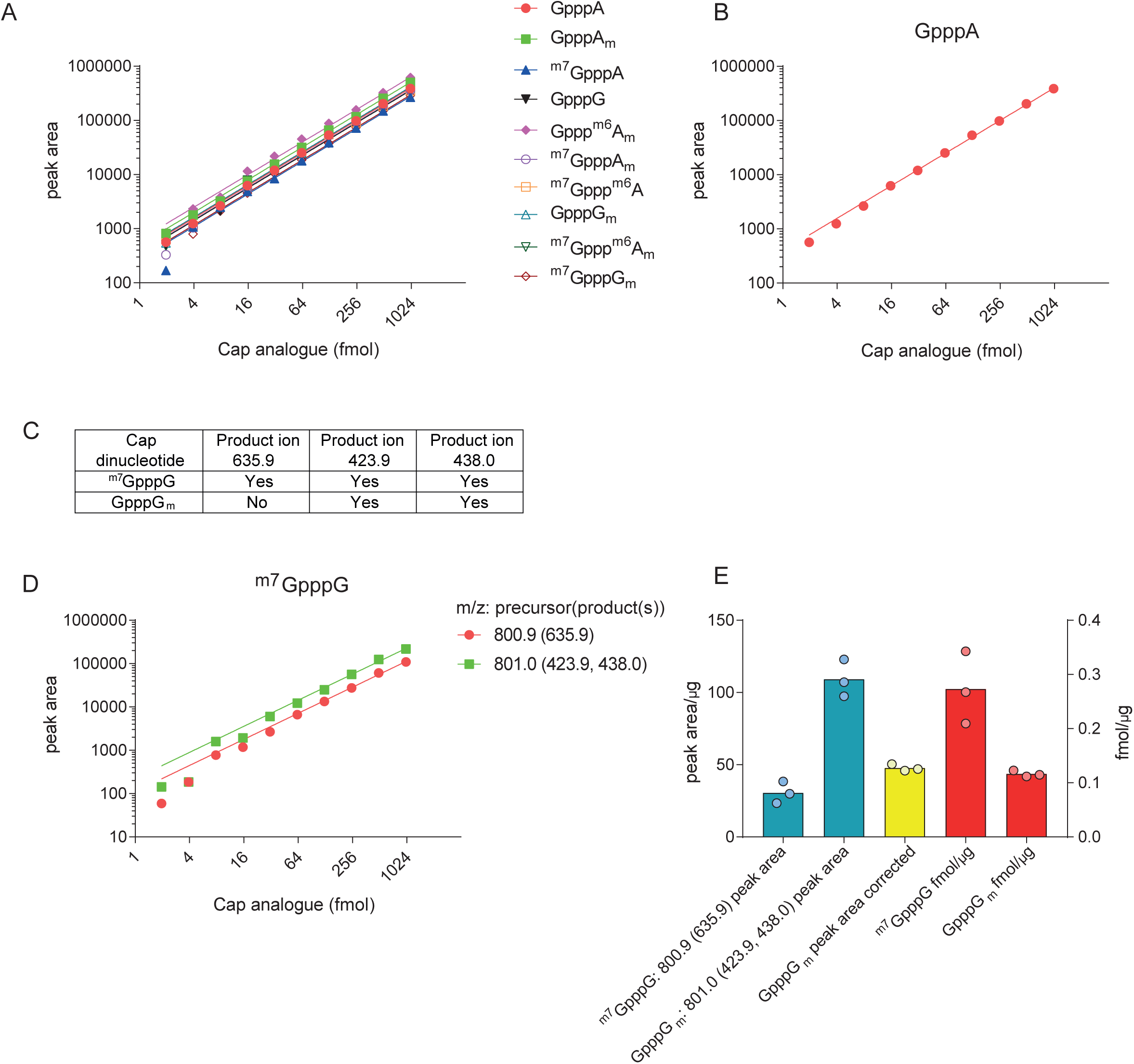
Detection of cap dinucleotides in serial dilution series. A) Peak area measurements from a serial dilution of synthetic cap dinucleotides. Slopes from linear regression of these values were calculated to allow conversion between peak area and fmol (Table 3). B) Peak area measurements for GpppA from a serial dilution of synthetic cap dinucleotides, provided as an example. C) Table demonstrating overlap in product ions originating from ^m7^GpppG and GpppG_m_. D) Detection of ^m7^GpppG across a dilution series. ^m7^GpppG is detected with its unique *m/z* 800.9 -> 635.9 precursor -> product ion transition, but it also contributes to the *m/z* 801.0 -> 423.9 and 438.0 precursor -> product ion transitions shared with the isobaric dinucleotide GpppG_m_. The *m/z* 801 -> 423.9 and 438 transition ion current signals from ^m7^GpppG can be back-calculated from the *m/z* 800.9 -> 635.9 precursor -> product ion signal and subtracted from the total *m/z* 800.9 -> 635.9 precursor -> product ion signal to allow quantification of GpppG_m_. E) Compensation for shared ions between ^m7^GpppG and GpppG_m_. ^m7^GpppG and GpppG_m_ raw peak areas are shown and the GpppG_m_ peak area after correcting for ^m7^GpppG forming product ions at m/z 423.9 and 438. Fmols of cap per µg input mRNA are calculated using the linear fit between fmols and peak area (Table 3). Each point represents a biological replicate.

### LC-MS analysis of mRNA cap dinucleotides from liver

Having established the CAP-MAP protocol with synthetic cap dinucleotides we proceeded to analyse cellular mRNA cap structures. Digestion of cellular mRNA with nuclease P1 results in a mixture of cap dinucleotides, nucleotide monophosphates and the nuclease protein. Various strategies including weak anion exchange and precipitation protocols were trialled to enrich and concentrate cap dinucleotides, however, these attempts resulted in a substantial loss of sample (data not shown). We were also concerned that the different enrichment methods might selectively alter the composition of cap dinucelotides in a sample. Therefore, we analysed cellular cap dinucleotides by LC-MS with minimal processing by simply directly injecting the whole nuclease P1 digests. Since there was no purification of the cap nucleotides after digestion, there was a risk that the nuclease P1 might accumulate on the column, increasing the back pressure. However, the back pressure of the column was monitored over 100 runs and was stable at around 40 bar.

When we analysed the mRNA cap structures present in mouse liver, we were able to detect nine of the eleven cap dinucleotides assessed (Fig.5A). Importantly, we found that the absolute ion currents recorded for the ARCA internal standard were the same whether or not it was injected alone or with cellular mRNA samples, suggesting no detectable interference (ion suppression) from the cellular mRNA nuclease P1 digests. Since PGC columns resolve structurally related compounds, it is likely that non-cap nucleotides bind to the column, but elute differentially to cap nucleotides, explaining the minor impact on the detection of cap structures.

**Figure 5.**
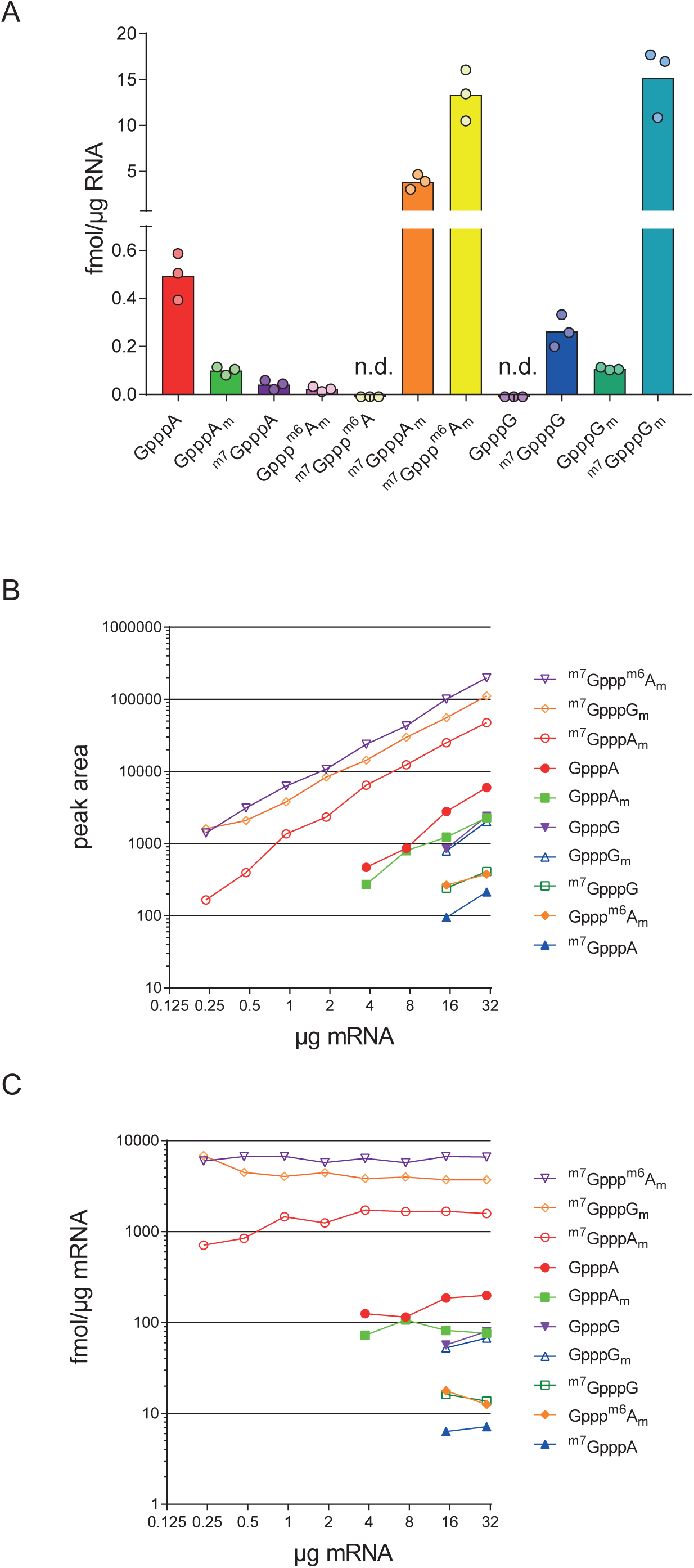
mRNA cap dinucleotides detected in mouse liver. A) Abundance of mRNA cap dinucleotides isolated from mouse liver. Each point indicates a biological replicate “n.d.”, indicates that a cap dinucleotide was not detected. B) Peak area measurements of different mRNA cap dinucleotides in a serial dilution of mouse liver mRNA. C) Calculated relative abundance (in fmol/µg mRNA) of different cap dinucleotides in a serial dilution of mouse mRNA.

The most common cap structures in liver mRNA were ^m7^GpppG_m_ (19 fmol/µg), ^m7^GpppA_m_ (4 fmol/µg), and ^m7^Gppp^m6^A_m_ (20 fmol/µg) (Fig.5A); these are the major cap variants previously elucidated by mass spectrometry and thin layer chromatography of radiolabelled nucleotides [1,29]. In line with recent observations that ^m7^Gppp^m6^A_m_ is an abundant cap structure, ^m7^Gppp^m6^A_m_ constituted 75% of the caps when A is the first transcribed nucleotide [10-12,43]. A number of incompletely methylated cap structures were detected including GpppA, GpppA_m_, Gppp^m6^A_m_, and GpppG_m,_ which lack cap guanosine N-7 methylation; and ^m7^GpppA, and ^m7^GpppG which lack ribose O-2 methylation of the first transcribed nucleotide.

To determine the minimum input of mRNA required to detect the different cap structures, we prepared a dilution series of liver mRNA. Ten cap dinucleotides, including in this experiment GpppG, were detected in 30 µg and in 15 µg mRNA, and the three most abundant caps were reproducibly detected in 250 ng mRNA (Fig.5B, C). The concentration (fmol/µg) of each cap dinucleotide was consistent across a range of inputs, when above the threshold of detection for that cap. The consistent detection of mRNA caps and ARCA internal standard across a range of input mRNA are further evidence of insignificant ion suppression from the cellular mRNA nuclease P1 digests.

In summary, we established CAP-MAP as a rapid, quantitative and relatively direct method for detecting mRNA cap dinucleotides.

### CAP-MAP comparison of mRNA caps from mammalian cells and tissues

To compare the relative cap abundancies in tissues, mRNA samples were prepared from liver, heart, brain and activated CD8 T cells. In each sample the most common cap dinucleotides were ^m7^GpppG_m_ and ^m7^Gppp^m6^A_m_ which were present at greater than 10 fmol/µg across all sources (Fig.6A). ^m7^GpppA_m_ was less common, but always present over 0.5 fmol/µg mRNA. A variety of less common cap dinucelotides were detected in the tissues and T cells; GpppA_m_, ^m7^GpppG, and GpppG_m_ were detected above 0.1 fmol/µg in all tissues; GpppA was present at ∼0.5 fmol/ µg in the heart and liver, but inconsistently detected in other tissues; ^m7^GpppA was present in low amounts in all samples except for one of the brain samples in which it was not detected; and Gppp^m6^A_m_ was very low in abundance and only consistently detected in liver (Fig.6B). Overall the minor cap variants, which lack either cap guanosine N-7 methylation or first nucleotide ribose O-2 methylation constituted around 2-5% of the cellular caps.

**Figure 6.**
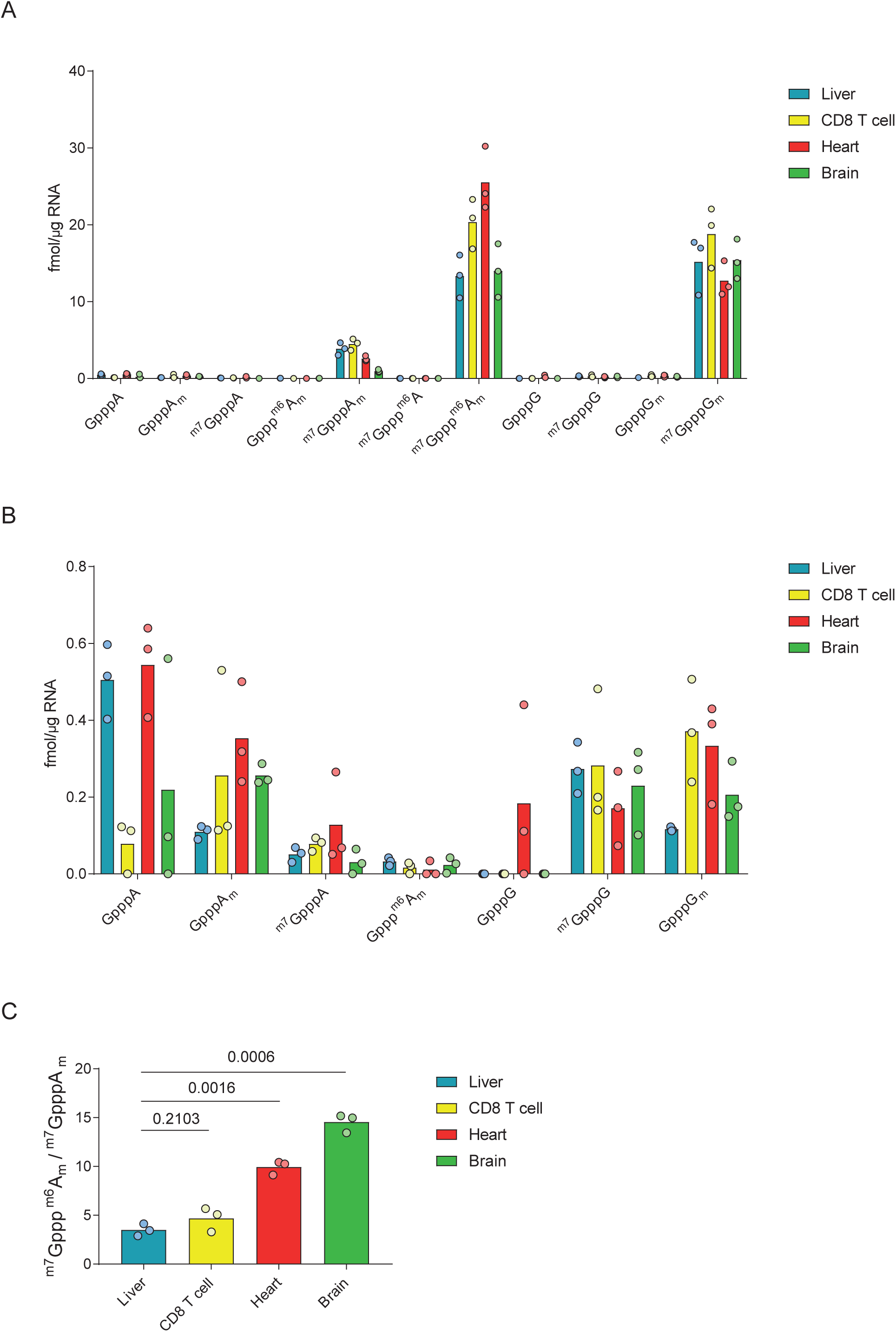
mRNA cap dinucleotides detected in mouse organs. A) Abundance of mRNA cap dinucleotides isolated from mouse liver, activated CD8 T cells, heart and brain. Each point indicates a biological replicate. B) Data from A presented to reveal the abundance of rarer mRNA cap dinucleotides (abundance <1 fmol/µg mRNA). C) Ratio of ^m7^Gppp^m6^A_m_ to ^m7^GpppA_m_ in mRNA from different sources.

Amongst the more common cap dinucleotides there were tissue-specific differences (Fig.6A, B). In most samples the abundance of caps with the first transcribed nucleotide A and G were similar, however in the heart there was more than twice as many caps with A as the first transcribed nucleotide compared to G. This likely reflects differences in the heart transcriptome, compared to other tissues analysed. The other major difference in cap structure abundance between tissues was the proportion of first nucleotide adenosine caps with the ^m6^A modification. All tissues had a greater concentration of ^m7^Gppp^m6^A_m_ than ^m7^GpppA_m_, but the ratio of these two cap structures varied considerably between tissues. Liver had the lowest ^m7^Gppp^m6^A_m_: ^m7^GpppA_m_ ratio of 3.5. Brain, at the other extreme, had a ^m7^Gppp^m6^A_m_: ^m7^GpppA_m_ ratio of 14.6 (Fig.6 C). These results indicated tissue-specific regulation of the ^m6^A cap methyltransferase, CAPAM, and/or the demethylase FTO. To demonstrate that our method can detect changes in the mRNA cap structure following perturbations in cap methyltransferase activity we reduced CAPAM expression by transfection of siRNA in HeLa cells. CAPAM knockdown was confirmed by western blotting (Fig.7A). As expected, CAPAM knockdown results in an increase in ^m7^GpppA_m_ and a decrease in ^m7^Gppp^m6^A_m_ (Fig.7B). In the control siRNA transfected HeLa cells there is about 9-fold more ^m7^Gppp^m6^A_m_ than ^m7^GpppA_m_, whereas this ratio drops to about 1.2-fold when CAPAM expression is repressed (Fig.7C). Notably the amount of ^m7^GpppG_m_ which is not a CAPAM substrate or product is very similar between the control and CAPAM siRNA treated cells. These findings demonstrate that short term regulation of a cap methyltransferase causes sufficient changes in mRNA structures to be detectable by CAP-MAP.

**Figure 7.**
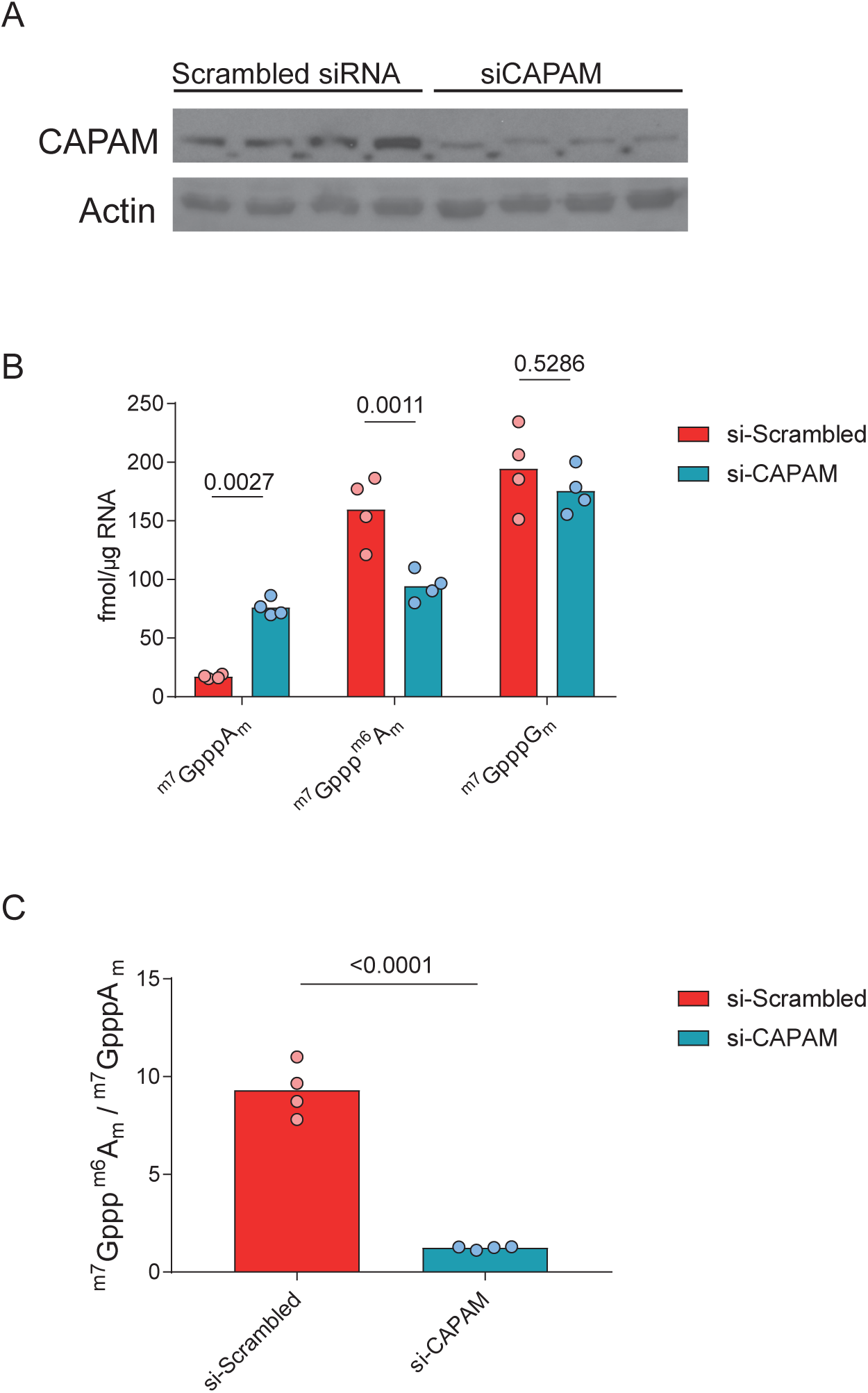
Impact of CAPAM knock-down on cap dinucleotide abundance. HeLa cells were treated with scrambled siRNA or siCAPAM for three days. A) Western blot of CAPAM and Actin (from re-probing) in siRNA-treated HeLa cells. B) Abundance of mRNA cap dinucleotides in siRNA-treated HeLa cells. Samples were compared by an ANOVA, with Sidak’s post-test, adjusted p values are shown. C) Ratio of ^m7^Gppp^m6^A_m_ to ^m7^GpppA_m_ in mRNA from siRNA-treated HeLa cells. Samples were compared by a Student’s t test.

## Discussion

Here we report the development of CAP-MAP, a rapid, sensitive and relatively direct method of mRNA cap analysis. The benefit of CAP-MAP is that following oligo-dT enrichment of mRNA and digestion, the cap structures are ready for LC-MS (liquid chromatography-mass spectrometry) analysis. This simple preparation is possible due to the selectivity of the PGC column and unique triphosphate linked dinucleotide structure of the mRNA caps. The small number of preparative steps means that the relative proportion of cellular cap structures are likely to be faithfully maintained during the analysis.

From the cap structures which we investigated (those with first nucleotide G or A), the major cap structures, ^m7^GpppG_m_, ^m7^GpppA_m_ and ^m7^Gppp^m6^A_m_ were detected in 250 ng mRNA (oligo-dT purified) from tissue sources. Minor, undermethylated cap structures were detected in 15 µg mRNA. As reported by earlier studies, caps methylated on the N-7 position of the terminal guanosine and O-2 position the first transcribed nucleotide ribose were the most common across a range of tissue sources/cell types [1,16,19,20]. However, in 2-5% of oligo-dT purified transcripts we also detected a variety of incompletely methylated cap dinucleotide structures in which one or both methyl groups were absent. Demethylase activities which target the methyl groups on the N-7 position of the cap guanosine or the O-2 position of the first transcribed nucleotide ribose have not been isolated from mammalian cells and therefore the undermethylated structures are unlikely to be breakdown products. This implies that either a certain amount of mRNA is processed into polyadenylated transcripts in the absence of a completed cap structure and/or the cap is completely removed, recapped and then partially methylated [44]. These undermethylated caps were uncommon, typically making up less than 5% of the total mRNA caps, indicating either that they are efficiently converted to their methylated forms and/or that they are effectively removed by decapping enzymes including DXO and NUDT16 [45,46].

The sequence of cap methylations is of mechanistic interest when considering regulation of mRNA cap formation. The cap0, cap1, cap2 notation, abbreviating ^m7^GpppN, ^m7^GpppN_m_, ^m7^GpppN_m_N_m_, respectively, can imply that guanosine N-7 methylation occurs prior to first nucleotide ribose O-2 methylation, however, to our knowledge, data has not been presented to suggest an obligatory order to methylation in mammalian cells. We detected both GpppN_m_ and ^m7^GpppN cap structures in a variety of tissues, suggesting that RNMT and CMTR1 can independently methylate the RNA cap. This is consistent with *in vitro* assays investigating the activity of these enzymes on GpppG caps and ^m7^GpppG caps [47,48]. Notably CMTR1 binds directly to the RNAPII C-terminal domain whereas RNMT interacts indirectly with RNAPII predominantly via interactions with RNA and RNAPII-associated proteins, which does not imply an obligate order of action [49,50].

CAPAM/PCIF1, the enzyme catalysing first transcribed nucleotide ^m6^A methylation has recently been identified and characterised [9-12]. ^m6^A methylation is the only cap methylation that has been demonstrated to be reversible leading to an interest in whether it may coordinate signalling events with translation or transcript stability [26]. Consistent with recent estimates of the abundance of the ^m7^Gppp^m6^A_m_ cap, we found it to be more abundant than ^m7^GpppA_m_ in all tissues [9,12,51,52]. However, the ratio of ^m7^Gppp^m6^A_m_ to ^m7^GpppA_m_ varied between tissues with higher ^m7^Gppp^m6^A_m_ in heart and brain, and lower ^m7^Gppp^m6^A_m_ in liver and CD8 T cells. This indicates that adenosine N-6 methylation by CAPAM or demethylation by FTO is differentially regulated across these tissues. CAP-MAP was able to detect a substantial decrease in ^m7^Gppp^m6^A_m_ levels and increase in ^m7^GpppA_m_ in HeLa cells following short term knock down of CAPAM illustrating its potential to investigate the effects of CAPAM and FTO regulation on cellular cap structures.

Recently an alternative LC-MS method for detecting mRNA caps was reported [53]. In addition to dinucleotide caps, a variety of rare non-canonical metabolite mRNA caps were detected in mammalian cells including NAD-caps, which have only recently been described in mammalian cells and FAD-caps, UDP-Gluc-caps and UDP-GlucNAc-caps which were not previously known to exist in mammals [54]. The physiological functions of metabolite-caps in mammalian cells remain uncertain and this is an active area of research [55]. In mouse liver and the human CCRF-SB cell line mRNA, Wang *et al*. also detected substantial amounts of ^m7^Gppp^m6^A, equivalent in concentration to ^m7^GpppA_m_. This was an unexpected finding because in previous analyses of HEK293T and MEL624 cells, this cap was not detected [9,12,51,52]. CAPAM was determined to have a 7.6-fold preference for ribose O-2 methylated caps over ribose unmethylated caps, a selectivity which would permit the formation of ^m7^Gppp^m6^A *in vivo* [10]. In CAP-MAP, we readily detected synthetic ^m7^Gppp^m6^A, but did not detect this cap dinucleotide in liver, nor any other mouse tissues. This could reflect either biological differences in the mRNA cap structures present in our samples, or differences in our separation and detection of ^m7^Gppp^m6^A. Notably we did detect other cap dinucleotides lacking O-2 methylation (discussed above), but these were minor variants.

To conclude, as it is becoming more apparent that mRNA cap methylation is a regulated process, a simple and sensitive method for quantitating the mRNA cap structures present in cells is required. Here we present CAP-MAP, a rapid and direct method for mRNA cap analysis by LC-MS. In CAP-MAP, major cap dinucleotide variants can be detected and quantified with low input mRNA (500ng) which can rapidly be recovered from cultured cells or tissue samples, and minor cap dinucleotides can be detected using a larger quantity of mRNA.

## Materials and Methods

### Cap dinucleotide standards

Synthetic cap dinucleotide standards were either bought from New England Biolabs or synthesised. A list of the cap dinucleotides and their sources is available in table 1. Syntheses of dinucleotide cap analogues GpppG_m_, GpppA_m,_ Gppp^m6^A_m_, ^m7^GpppG_m_, ^m7^GpppA_m,_ ^m7^Gppp^m6^A, ^m7^Gppp^m6^A_m_, were performed from GDP or m^7^GDP and imidazolide derivatives of 2’-O-methylguanosine 5’-O-monophosphate (m^2’-O^GMP), 2’-O-methyladenosine 5’-O-monophosphate (m^2’-O^AMP) and 6,2’-O-dimethyladenosine 5’-O-monophosphate (m_2_^6,2’-O^AMP), respectively, according to the methods reported previously [56,57]. The mononuclotides substrates (m^2’-O^GMP, m^2’-O^AMP, m^6^AMP, and m_2_^6,2’-O^AMP) were prepared from the appropriate nuclosides (m^2’-O^Guo, m^2’-O^Ado, m^6^Ado and m_2_^6,2’-O^Ado) by 5’-O-phosphorylation according to the procedures described earlier [56,58]. The intermediate 6-methyladenosine (m^6^Ado) was prepared from 6-chloropurine riboside (Aldrich) according to Johnson et al. (1958)[59]. 2’-O-Methylated intermediates m^2’-O^Guo, m^2’-O^Ado, m_2_^6,2’-O^Ado and were synthesised using methods described by Robins et al. (1974)[60].

### Cap dinucleotide detection and relative quantification by LC-MS

Eleven different cap dinucleotide standards were used to optimise the LC-MS method for cap detection and quantification. Cap nucleotides levels were measured using a TSQ Quantiva mass spectrometer interfaced with Ultimate 3000 Liquid Chromatography system (ThermoScientific), equipped with a porous graphitic carbon column (HyperCarb 30×1mm ID 3μm; Part No: C-35003-031030, Thermo-Scientific). Mobile phase buffer A consisted of 0.3% (v/v) formic acid adjusted to pH 9.15 with ammonia prior to a 1/10 dilution. Mobile phase buffer B was 80% (v/v) acetonitrile. The column was maintained at a controlled temperature of 45°C and was equilibrated with 18% buffer B for 9 minutes at a constant flow rate of 0.04 mL/min. Aliquots of 14 µL of each sample were loaded onto the column and compounds were eluted from the column with a linear gradient of 18-21% Buffer B over 2 mins, 21-40% Buffer B over 2 mins, 40-60% Buffer B over 10 mins, the column was then washed for 4 min in 100% Buffer B before equilibration in 18 % Buffer B for 9 mins. Eluents were sprayed into the TSQ Quantiva using Ion Max NG ion source with the ion transfer tube temperature at 350°C and vaporizer temperature at 60°C. The TSQ Quantiva was run in negative mode with a spray voltage of 3500, Sheath gas 40 and Aux gas 10 and sweep gas 2. Levels of the 11 cap nucleotides were measured using multiple reaction monitoring mode (MRM) with optimised collision energies and radio frequencies previously determined by infusing pure compounds (Table 1). A chemically similar cap standard (ARCA (^m7^G_O-3m_pppG)) (NEB) was used as internal standard (Table 1). A standard curve was prepared using 3 to 1000 fmole of all 11 cap nucleotides with 250 fmole internal ARCA standard spiked in to each sample. The standard curve was run prior to running the experimental samples using the same conditions, and was used to calculate the relative amount of the 11 cap nucleotides in each sample. One blank was run between each sample to eliminate carryover.

### Preparing RNA from tissues

Mice were either C57Bl/6J mice or control mice from our colonies that are kept on a C57Bl/6J background. Mice were housed in the University of Dundee transgenic facility. Livers, brains and hearts were snap frozen in liquid nitrogen, ground in a pestle and mortar and lysed in TRI Reagent (Sigma). RNA was extracted according to the manufacturer’s protocol including the optional centrifugation before the addition of chloroform to remove insoluble material. In addition, a second phenol chloroform extraction was performed on the aqueous phase from the initial separation in TRI Reagent (ThermoFisher). RNA was dissolved in Hypersolv water (VWR) and quantified using a nanodrop (ThermoFisher).

### Preparing RNA from cell cultures

Cells were cultured in 5% CO_2,_ 37ºC. CD8 T cells were purified from mouse lymph nodes using an easySep CD8 kit (Stemcell technologies) and activated on plate bound anti-CD3 (clone 145-2C11, Biolegend) and anti-CD28 (Clone 37.51, Biolegend) antibodies in RPMI medium (Gibco/Thermofisher) with 20ng/ml IL-2 (Novartis), 10% FCS (Gibco/Thermofisher), 50mM 2-ME (Sigma) and PenStrep (Thermofisher) for five days. Hela Cells were grown in DMEM (Gibco/Thermofisher) with 10% FCS and 5U/ml Penicillin-Streptomycin (ThermoFisher). PCIF1-directed siRNA smartpool or scrambled siRNA (Dharmacon) was transfected using a Neon transfection system (Thermofisher) using 1005V X 35ms X2. CAPAM knockdown was confirmed by western blotting with anti-PCIF1 (ab205016, Abcam) and anti-actin (ab3280, Abcam) antibodies. RNA was prepared from cells as per the tissue samples, with the exception that cells were directly lysed in Tri Reagent rather than frozen and ground.

### Preparing mRNA

For large scale mRNA extractions, polyadenylated mRNA was enriched using 100µl oligo-dT agarose beads (NEB) per mg RNA. RNA was bound to the beads in binding buffer (1M NH_4_OAc, 2mM EDTA) for 5 minutes rotating at room temperature and washed once. RNA was eluted in room temperature water, filtered in SpinX (Corning) tubes to remove any beads, then precipitated in 2.5M NH_4_OAc and isopropanol, centrifuged at 14K x 30mins and pellets washed in 75% Ethanol.

To extract mRNA from HeLa cell samples an mRNA Direct kit (ThermoFisher) was used. 40µl magnetic oligo-dT beads were used per sample, but with the binding buffer and incubation times indicated above. mRNA was digested with 9.5 units of Nuclease P1 (Sigma) in 20mM NH_4_OAC pH 5.3 for 3 hours at 37ºC. 250 fmols of ARCA was added, then LC-MS carried out as described above.

### Analysis

Peak areas were determined for each transition for each cap dinucleotide. Where there was more than one transition peak on the chromatogram, the correct peak was selected by its retention time relative to the ARCA internal standard and presence of product ions in the ratio indicated by the synthetic standards. The abundance of the various caps from mRNA were calculated by comparison with the dilution series of cap standards. Conversion factors are listed in Table 3. Statistical analysis and plots were produced in Prism 5 (GraphPad Software).

## Supplemental Figure Legends

**Supplemental figure 1: Representative chromatograms from the titration of liver mRNA from figure 5 B and C.**

30µg sample shown. Red arrows indicate the peak corresponding to the cap dinucleotide indicated.

**Supplemental figure 2: Representative chromatograms from the analysis of liver mRNA from figures 5 A and 6 A, B and C.**

Red arrows indicate the peak corresponding to the cap dinucleotide indicated.

**Supplemental figure 3: Representative chromatograms from the analysis of CD8 T cell mRNA from figure 6 A, B and C.**

Red arrows indicate the peak corresponding to the cap dinucleotide indicated.

**Supplemental figure 4: Representative chromatograms from the analysis of heart mRNA from figure 6 A, B and C.**

Red arrows indicate the peak corresponding to the cap dinucleotide indicated. These samples were run on a different PGC column to the other samples (which later stopped working), we found that there were column to column differences in the retention times, but the order of elution was maintained.

**Supplemental figure 5: Representative chromatograms from the analysis of brain mRNA from figure 6 A, B and C.**

Red arrows indicate the peak corresponding to the cap dinucleotide indicated.

**Supplemental figure 6: Representative chromatograms from the analysis of HeLa mRNA from figure 7 B and C.**

HeLa cells were treated with scrambled siRNA (A) or CAPAM siRNA (B). RT: retention time and A: peak area, are shown.

## Acknowledgments

We would like to thanks members of the Cowling lab and Fingerprints proteomics facility for useful discussions.

