## Supplemental Figures for "CAP-MAP: Cap Analysis Protocol with Minimal Analyte Processing, a rapid and sensitive approach to analysing mRNA cap structures"

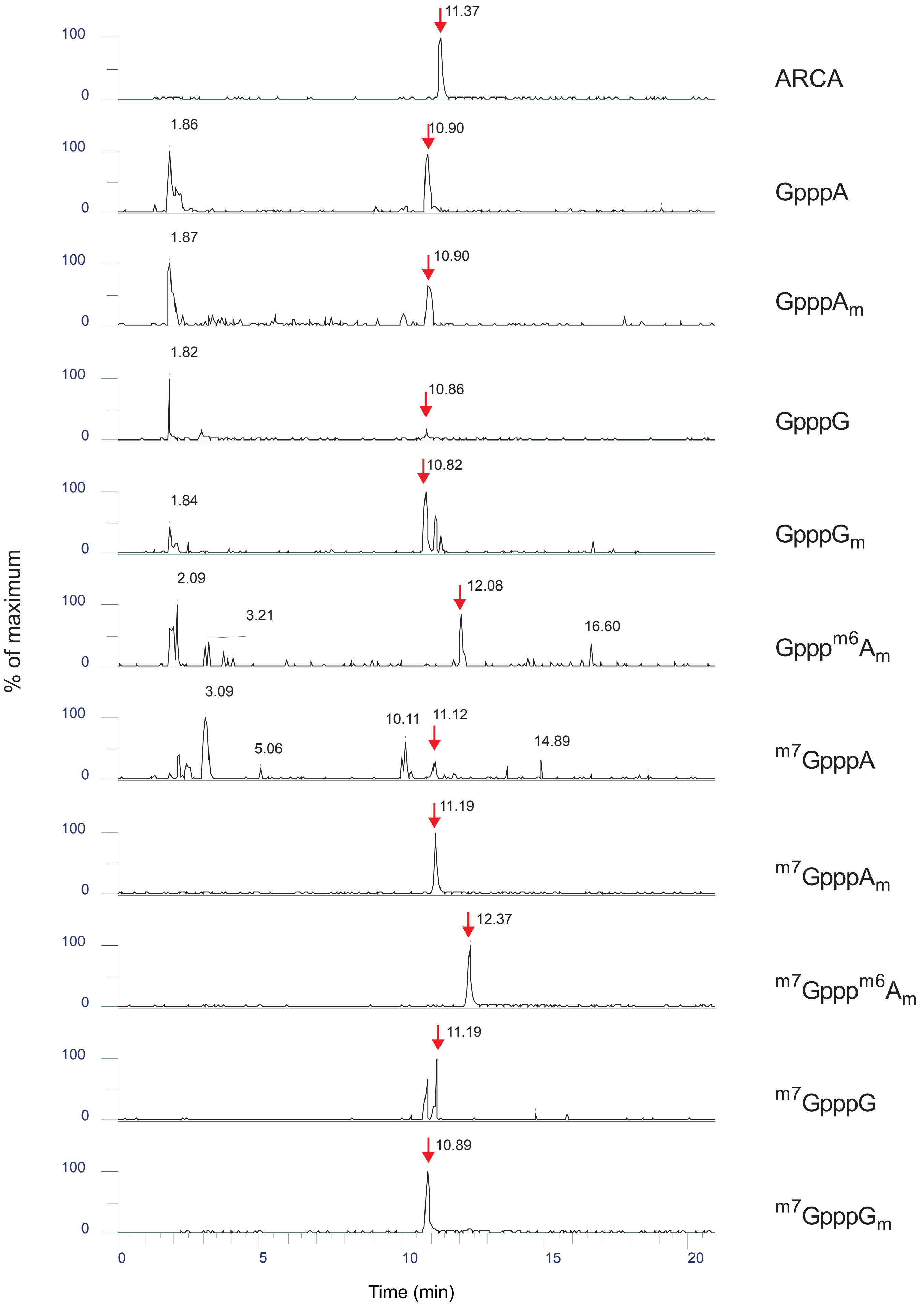


Supplemental figure 1

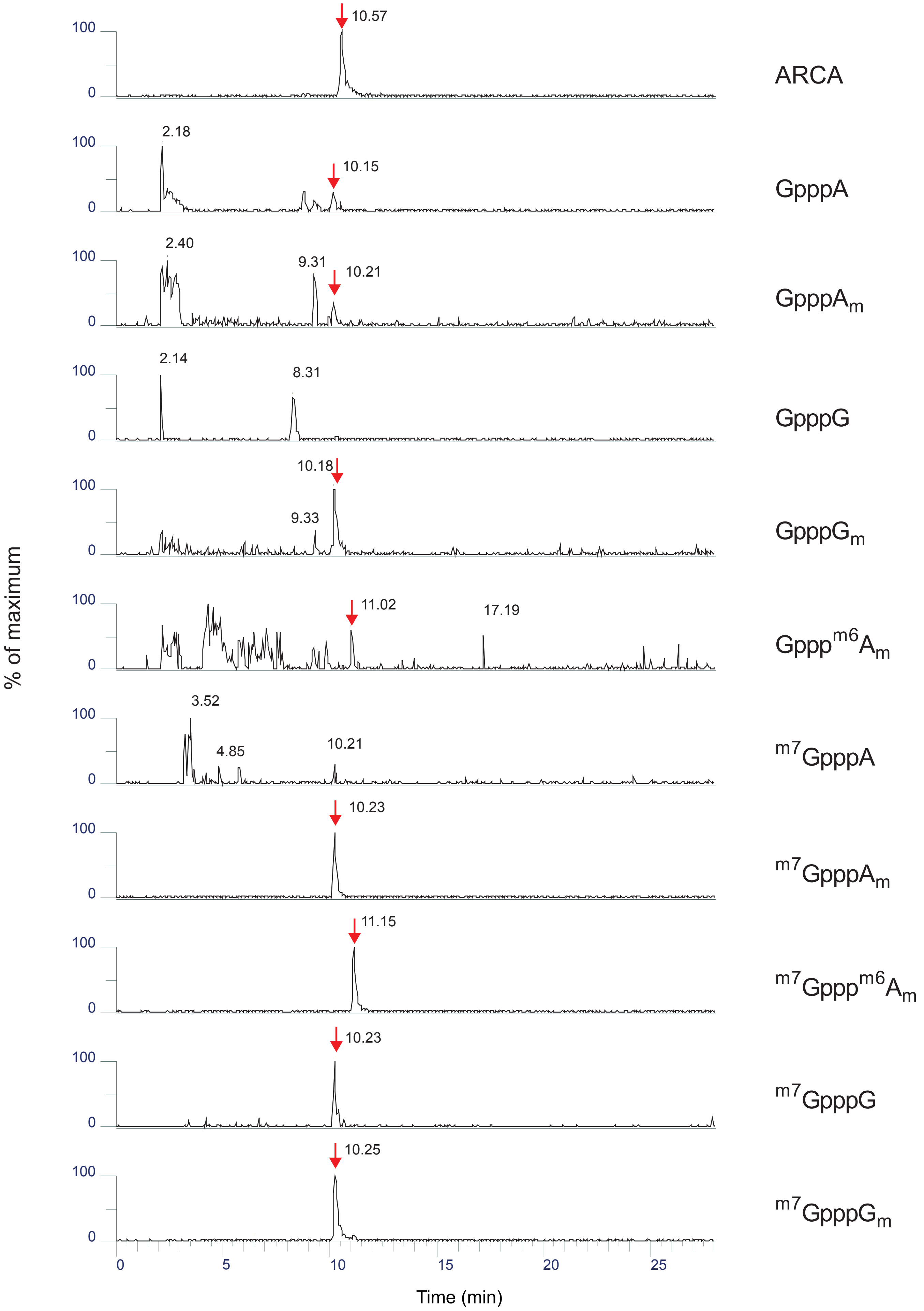


Supplemental figure 2

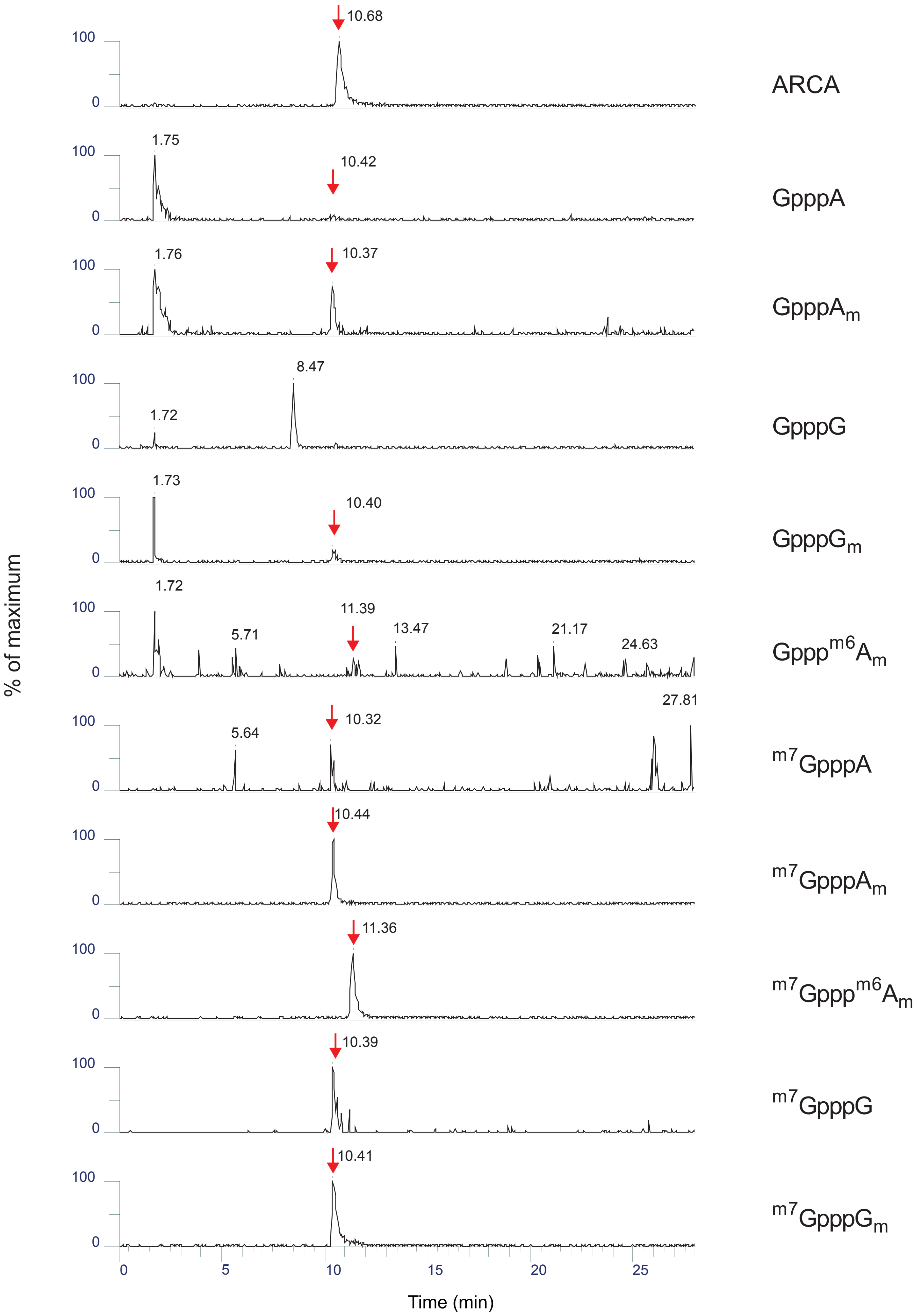


Supplemental figure 3

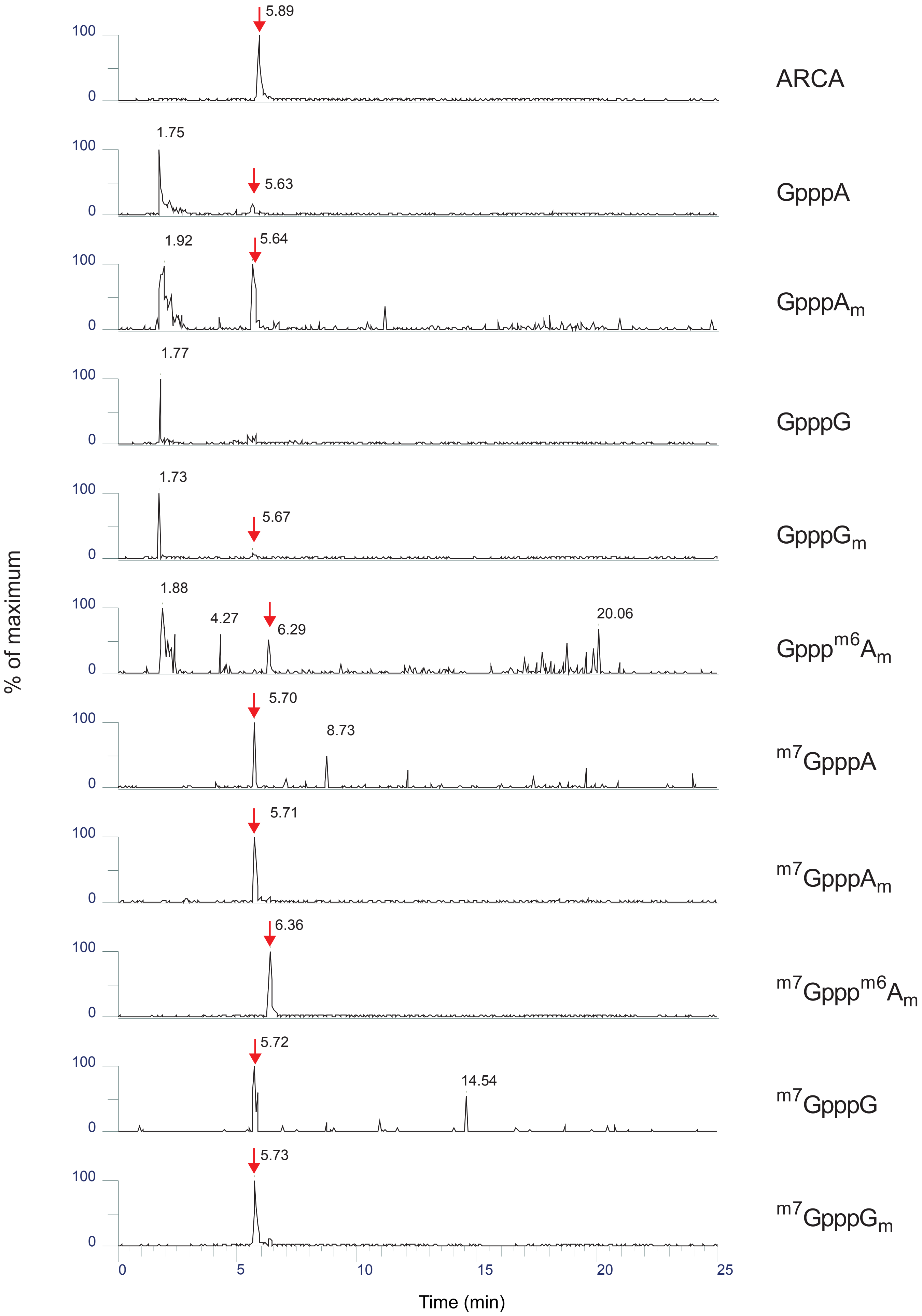


Supplemental figure 4

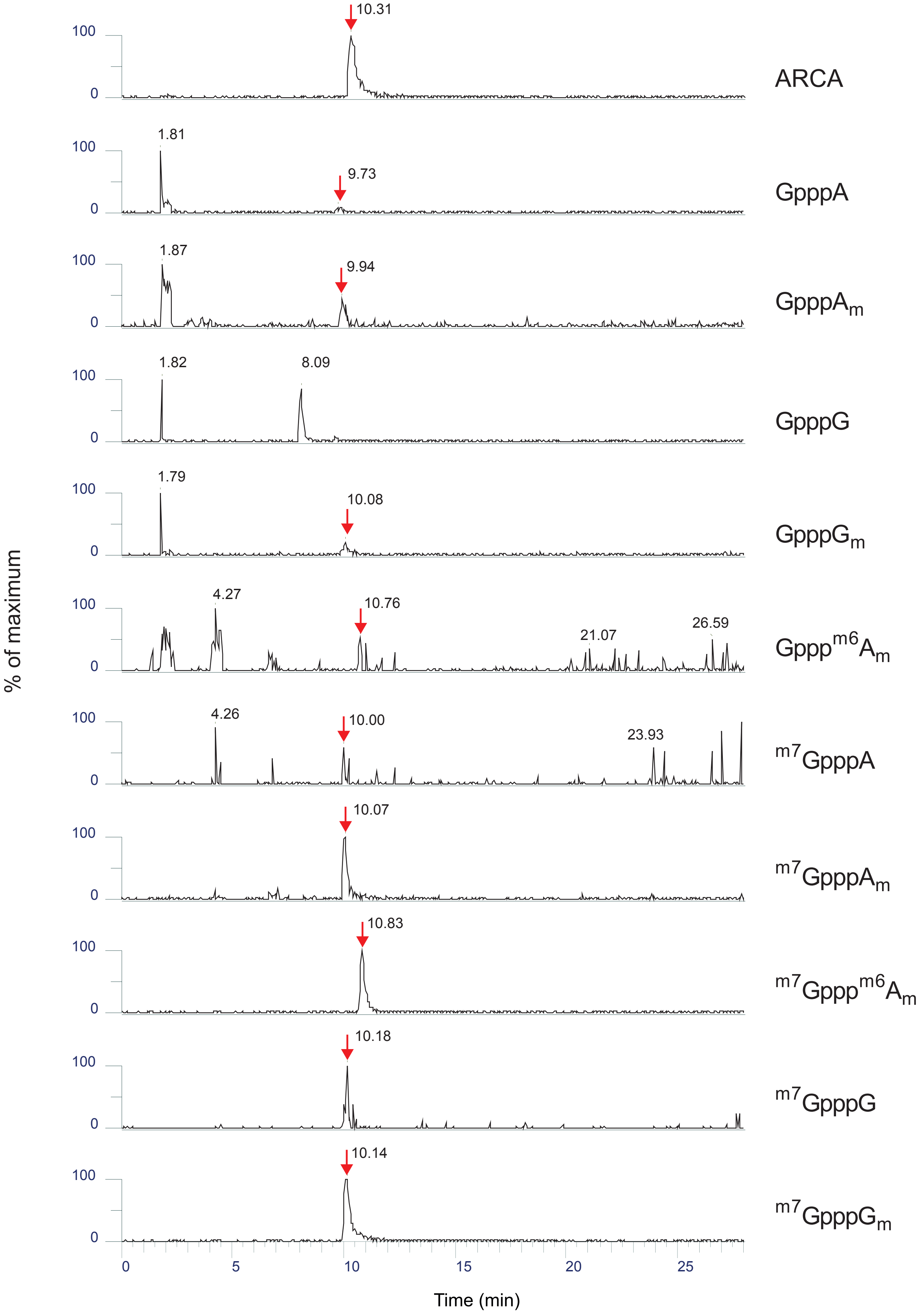


Supplemental figure 5

A

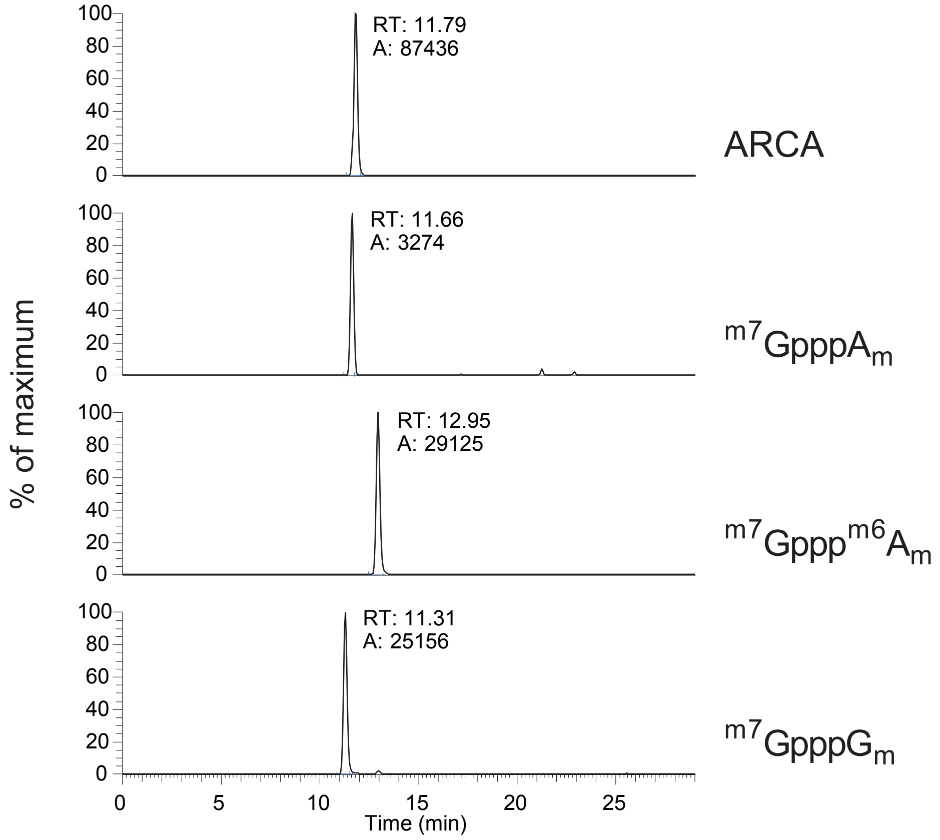


B

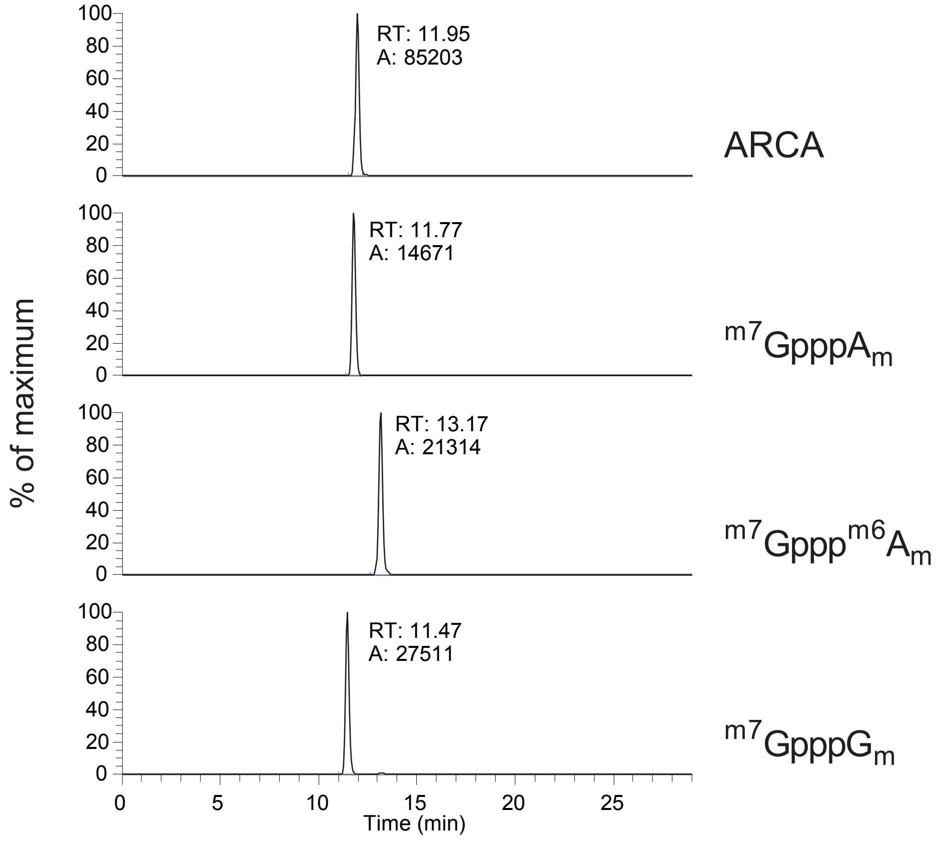


Supplemental figure 6
